# Intrinsic defects in lymph node stromal cells underpin poor germinal center responses during aging

**DOI:** 10.1101/2020.05.07.082255

**Authors:** Alice E Denton, Alyssa Silva-Cayetano, James Dooley, Danika L Hill, Edward J Carr, Philippe A Robert, Michael Meyer-Hermann, Adrian Liston, Michelle A Linterman

## Abstract

The failure to generate enduring humoral immunity after vaccination is a hallmark of advancing age. This can be attributed to a reduction in the germinal center response, which generates long-lived antibody-secreting cells that provide protection against (re)infection. Despite intensive investigation into the effect of age on the lymphoid compartment, the primary cellular defect that causes impaired germinal centers in aging has not been identified. Herein we demonstrate that aging reduces the capacity of germinal center-associated stromal cells to respond to vaccination. Heterochronic parabiosis and mathematical modeling demonstrate that a poor stromal cell response limits the size of the germinal center. This study reveals that age-associated defects in stromal cells are a significant barrier to efficacious vaccine responses in older individuals.

## Introduction

While the extension of human life expectancy is a great achievement of the modern era, it creates a challenge for science: to facilitate healthy aging. Older individuals are at particular risk of infection, and this is in part caused by decreased functionality of the immune system^1^. Vaccination is an effective preventative strategy for limiting the negative health consequences from infection, however older persons often do not generate long-lasting protective immunity after vaccination^2, 3, 4, 5^. Therefore, an understanding of why older individuals do not respond well to vaccines is key to developing the next generation of immunization strategies that are effective in older persons.

The generation of lasting protective immunity requires the production of long-lived plasma cells that secrete high affinity class-switched antibodies^6^. Plasma cells are the ultimate output of the germinal center (GC), a specialized immune response that forms in secondary lymphoid organs^7^. The GC reaction requires the concerted efforts of many different cell types that must be coordinated in time and space: B cells, T follicular helper (Tfh) cells, dendritic cells and macrophages are brought together upon a network of specialized stromal cells that provide migratory cues that direct cells to, and within, the GC^8^. The GC is a polarized structure with two functionally distinct compartments known as light and dark zones^9, 10^. In the dark zone, GC B cells proliferate and the genes encoding the B cell receptor undergo somatic hypermutation. Because of the random nature of somatic hypermutation, selection must occur to ensure this process has not negatively impacted on the function or specificity of the B cell receptor, as well as increasing receptor affinity. To do this, GC B cells move to the light zone where they can receive help from a specialized subset of CD4^+^ T cells, Tfh cells^11^. GC B cells that received help from Tfh cells then migrate back to the dark zone and either undergo further rounds of proliferation and mutation or differentiate into memory B cells or long-lived antibody-secreting cells that exit the GC^7^.

Stromal cells are critical to lymph node structure, dictating the T and B cell areas and directly regulating immune cell homeostasis and adaptive immune responses^12, 13, 14^. Stromal cells form the physical scaffold upon which immune cells migrate and produce molecules that dictate the migration and survival of immune cells^15, 16, 17^. There is significant diversity in the types of lymph node stromal cells^18^; for the GC, two key stromal cells are known to dictate its structure and function. The light zone is supported by follicular dendritic cells (FDCs), which are essential for the GC^19^ as they produce CXCL13 and act as a depot for antigen^20^. The dark zone is populated by CXCL12-abundant reticular cells whose functions are poorly defined outside of their role as a migratory nexus^10, 21, 22^. A third cell type, marginal reticular cells (MRCs), may also play a role in GC formation as they may contribute to antigen transport^23, 24^ and can act as a progenitor for *de novo* FDC differentiation^25^.

A reduction in the size, quality and output of the GC in older individuals is observed in a number of infection and vaccination settings^26, 27, 28, 29^. This has been linked to T cell-intrinsic mechanisms^30, 31, 32, 33^, and changes in the microenvironment^34, 35, 36, 37, 38^. Age reduces the ability of FDCs to capture and retain antigen^27, 39, 40^, suggesting an impaired functionality in aging that may underpin poor GC formation. However, apart from changes in antigen capture by FDC, we do not know how GC stromal cells are influenced by age and what contribution any cell-intrinsic changes make to the defective GC reaction. We show here that aged FDCs and MRCs respond poorly to immunization, at a cellular and molecular level. MRCs are particularly affected, with age causing a reduction in their proliferative response to immunization that limits the expansion of the FDC network through *de novo* differentiation. We use mathematical modeling and heterochronic parabiosis to demonstrate that the poor responsiveness of stromal cells directly impairs the capacity of B cells to participate in a GC, independent of B cell age. In sum, this study identifies the impact of age on the transcriptional and cellular responses of MRCs and FDCs to immunization and emphasizes that these defects in stromal cell responses underpin the poor GC response in aged individuals.

## Results

### Aging impairs GC and FDC expansion in response to immunization

In order to understand how stromal cells are affected by aging, and how these changes impact on the generation of GC responses to immunization, we subcutaneously immunized adult (8-12 week old) and aged (>95 week old) Balb/c mice. The ki67^+^Bcl6^+^B220^+^ GC B cell response (Fig 1a) was reduced in both proportion (Fig 1b) and absolute number (Fig 1c) consistent with previous studies^26, 27, 34^. The reduction in GC size correlated with a reduced FDC network, where CD21/35^+^ FDCs (Fig 1d) were reduced in proportion (Fig 1e) and absolute number (Fig 1f). To determine whether these results were consistent between mice of different genetic backgrounds, we immunized C57BL/6 mice and determined the GC B cell response seven days later (Fig 1g). Both the proportion (Fig 1h) and absolute number (Fig 1i) of GC B cells was reduced in aged mice. As observed in Balb/c mice, we also found a correlation between reduced GC magnitude and reduced FDC expansion, where FDCs were significantly reduced when measured either by proportion (Fig 1j, k) or in absolute number (Fig 1l) in C57BL/6 mice. This demonstrates that ageing negatively influences the response of GC-associated stromal cells to immunization.

**Figure 1.**
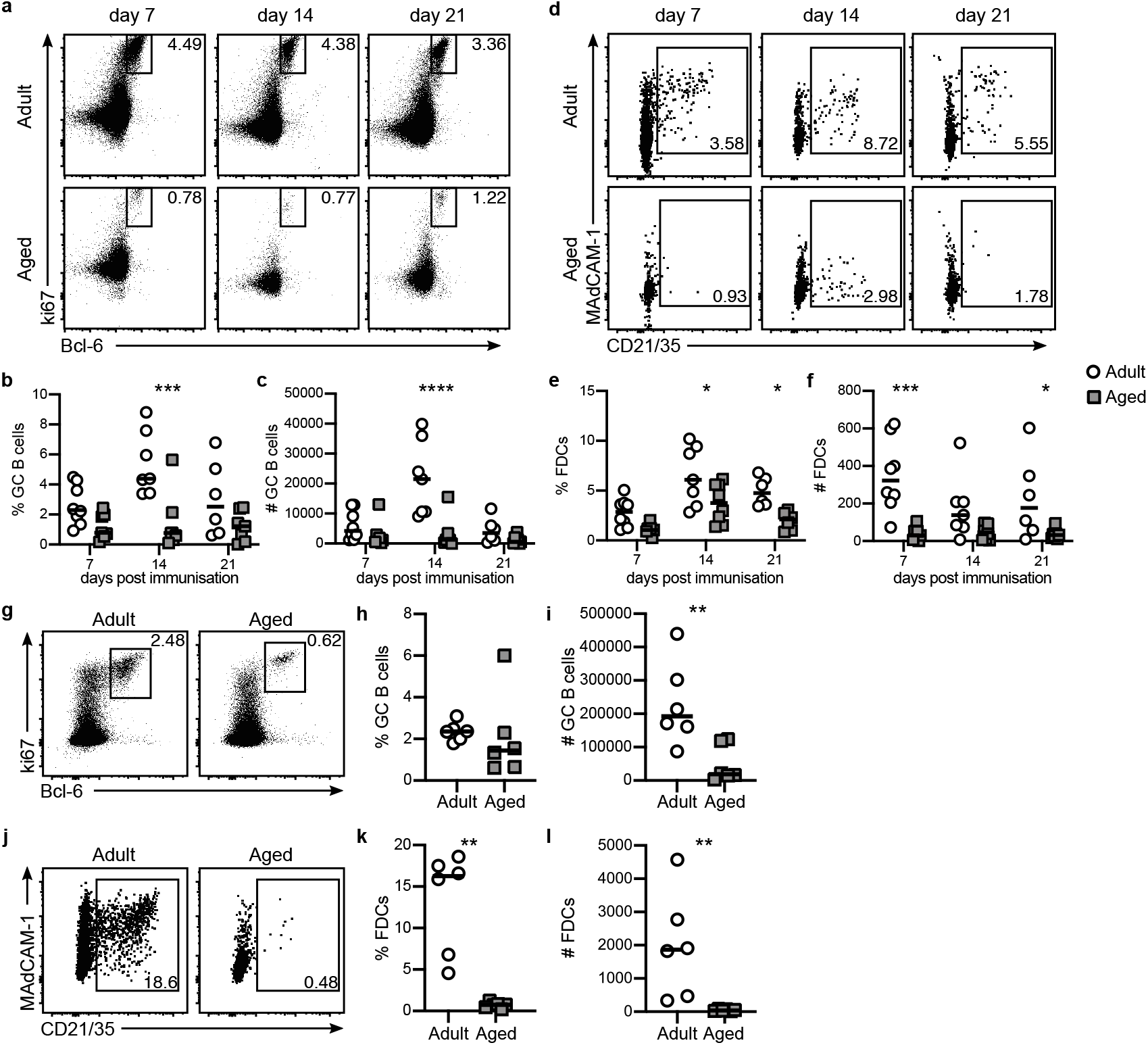
Aging diminishes the GC and FDC response to immunization. Adult (8-12 week old) and aged (>98 week old) mice were immunized subcutaneously with 50μg NP-KLH emulsified in Alum and the stromal and immune cell responses were determined. (a-f) GC B cell and FDC responses were determined in Balb/c mice seven, 14 and 21 days after immunization. (a) Representative gating of Bcl-6^+^ki67^+^ GC B cells. (b-c) The proportion (b) and absolute number (c) of GC B cells at different times after immunization. (d-f) FDC responses were determined for the same timepoints as in (a-c). (d) Representative gating of and FDCs (CD21/35^+^) amongst CD31^-^CD45^-^Pdpn^+^ stromal cells. (e-f) The proportion (e) and absolute number (f) of FDCs. (g-l) GC B cell and FDC responses were determined in C57BL/6 mice seven days after immunization. GC B cells were gated as Bcl-6^+^ki67^+^ (g), and the proportion (h) and absolute number (i) of GC B cells was determined. FDCs were gated based on CD21/35 staining amongst CD31^-^CD45^-^Pdpn^+^ stromal cells (j) and the proportion (k) and absolute number (l) of FDCs was determined. Each symbol represents a biological replicate and lines indicate the median. Data are representative of at least two independent experiments with 6-8 mice per group. Statistical significance was determined using a Mann-Whitney rank test. * p<0.05, **p<0.01, ***p<0.001, ****p<0.0001.

### Aging impairs the transcriptional response of FDC to immunization

Having established that the number of FDCs in a mature GC is reduced in age, we wanted to understand if there were molecular differences in how FDCs respond to immunization, and how this is altered with advancing age. To address these questions, we performed RNA sequencing on sort-purified FDCs (CD21/35^+^Pdpn^+^CD31^-^CD45^-^) isolated from lymph nodes of adult and aged mice both prior to and two days after immunization. We elected to study FDCs isolated early after immunization in order to understand how the immunization directly affects the transcriptome of FDCs, rather than capturing changes that are secondary to the initiation of adaptive immune responses, and also minimizing the risk of confounding due to differences in FDC number. FDC identity was confirmed by the lack of immune and endothelial cell-related gene expression (Extended data Fig 1a) and high expression of known FDC-associated genes^18^, including *Cxcl13*, *Cxcl12*, *Cr2*, *Fcgr2b* and *Pthlh* (Extended data Fig 1b, c); the reduction in *Fcgr2b* in aged FDCs is consistent with previous studies^40^. Principal component analysis (PCA) revealed that there was little structure to the data that could be explained by age or immunization (Fig 2a). Supervised analysis of the transcriptome of FDCs revealed minor gene expression changes driven by age or immunization. In resting FDCs, age significantly affected the expression of 105 genes (Fig 2b, Extended data Fig 2a, b), and immunization significantly affected the expression of 110 genes in adult FDCs (Fig 2c, Extended data Fig 2d, e). Immunization of aged mice induced far fewer changes in FDCs than that observed in adult mice, where only 44 genes were differentially expressed (Fig 2d, Extended data Fig 2f, g). Very little overlap was observed in the immunization-induced gene expression changes between adult and aged FDCs (Fig 2e), suggesting that the transcriptional response to immunization in FDCs, whilst modest, was deregulated in the aged setting. Notwithstanding these findings, the small overall transcriptional change in FDCs following immunization suggested to us that early age-associated cell-intrinsic changes in FDCs themselves may not underpin the defect in stromal cells observed in aged mice, and prompts the hypothesis the defect may lie with their precursors.

**Figure 2.**
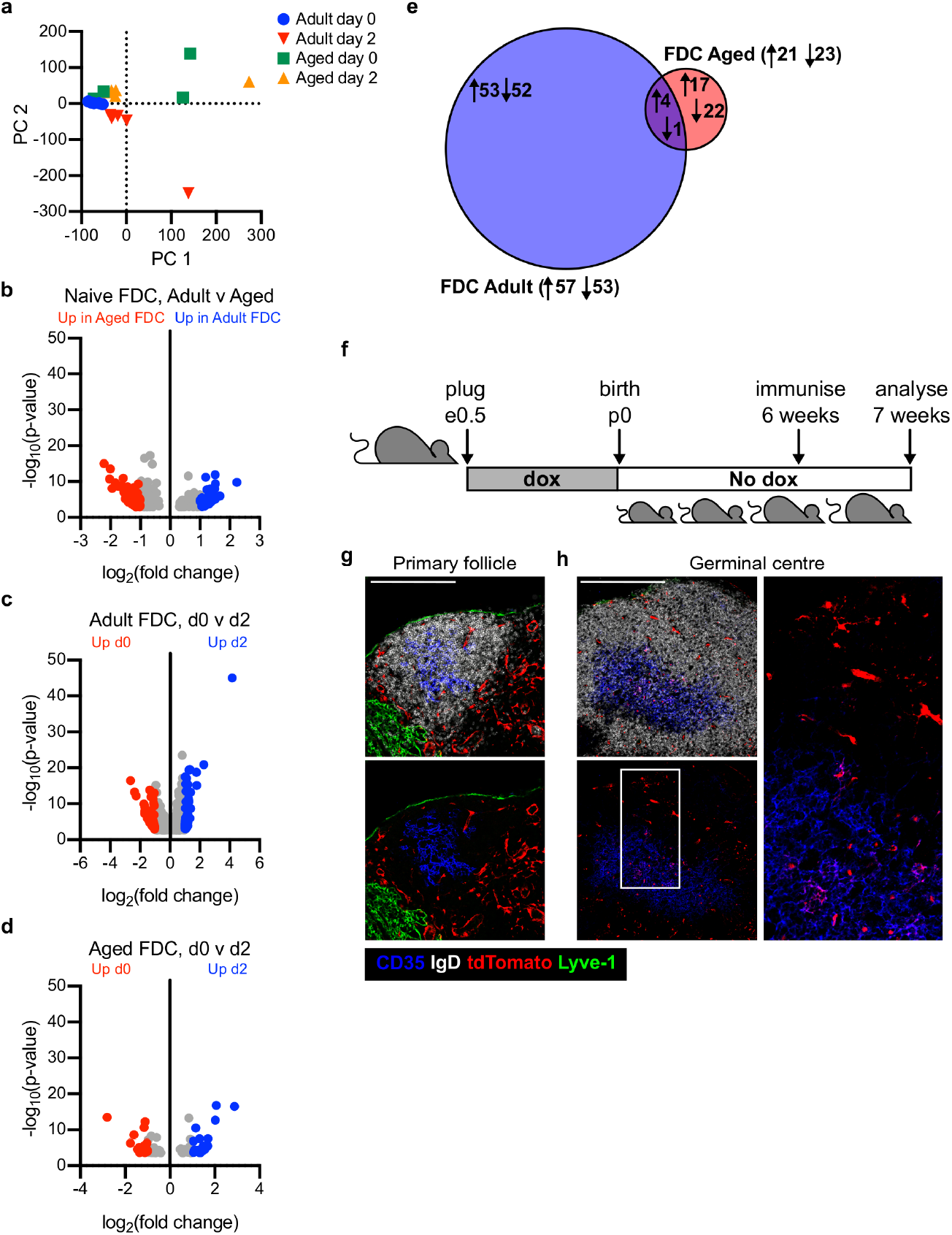
Transcriptional profiling of the FDC response to immunization in aging. (a-e) FDCs (CD21/35^+^Pdpn^+^CD31^-^CD45^-^) were isolated from adult (8-12 week old) and aged (>98 week old) unimmunized mice, or mice immunized two days prior with 50μg NP-KLH emulsified in Alum, and their transcriptomes sequenced. (a) PCA of all expressed genes in all FDC samples. (b) Volcano plots showing all genes that are differentially expressed (DESeq2) between adult and aged FDCs isolated from unimmunized mice. Genes that are enriched in aged (red) or adult (blue) samples are highlighted. (c, d) Volcano plots showing all genes that are differentially expressed (DESeq2) between naïve and immunized FDCs in adult (c) and aged (d) mice. Genes that are enriched in unimmunized (red) or immunized (blue) FDCs are highlighted. (e) Comparison of differential gene expression following immunization in FDCs isolated from adult or aged mice, where the overlap shows genes that change in both adult and aged FDCs. (f-h) *FAP*^tTA^::*TetO*^Cre^::*R26R*^tdTomato^ mice were administered doxycycline from prior to conception until birth then maintained in the absence of doxycycline. At six weeks of age, mice were immunized and tissues taken for analysis seven days later (f). Cryosections (20μm) were prepared and stained for IgD (white), Lyve-1 (green) and CD35 (blue), alongside tdTomato (red). (g) Lack of tdTomato labeling of FDCs in a primary B cell follicle in the non-draining lymph node. (h) tdTomato labeling of FDCs in a GC in the draining lymph node. Scale bars 200μm. Data in (a-f) are from five biological replicates except for the aged unimmunized FDC group, where four replicates were sequenced. Data in (f-h) are representative for four separate immunizations.

It has previously been suggested that MRCs can act as a precursor for *de novo* FDC differentiation in response to immunization^25^. To confirm this finding, we used a lineage-tracing model based on temporal control of fibroblast activation protein-alpha (FAP) expression, which we previously used to demonstrate that different lymph node stromal cells are developmentally related^41^. FAP-driven labeling of lymph node stromal cells was inhibited during embryogenesis by administering doxycycline from conception until birth (e0-p0). At six weeks of age mice were immunized and the labeling of FDCs in GCs was assessed seven days later (Fig 2f). In this system, most T cell zone stromal cells and some MRCs in primary B cell follicles were indelibly labeled with tdTomato (Fig 2g), consistent with our previous observations^41^. In GCs, which we defined by lack of IgD within an IgD^+^ follicle, tdTomato^+^ FDCs (Fig 2h) were observed, suggesting *de novo* FDC differentiation from non-FDCs, as FDCs do not actively express FAP^13, 41^. Because tdTomato^+^ FDCs were linked in a chain to the follicle edge (Fig 2h, inset), we conclude that, as previously described^25^, MRCs adjacent to the adult follicle can differentiate into FDCs following immunization. These data led us to investigate the effect of aging on the MRC response to immunization, as defects in this subset may explain the poor FDC response in aged mice.

### Aging significantly perturbs the MRC response to immunization

In order to understand the MRC response to immunization, we immunized adult (8-12 week old) and aged (>95 week old) mice and determined the number of MRCs, defined as MAdCAM-1^+^CD21/35^-^ amongst Pdpn^+^CD31^-^CD45^-^ stromal cells (Fig 3). In Balb/c mice (Fig 3a-c), we observed a trend towards a reduction in MRCs as a proportion of CD45^-^CD31^-^Pdpn^+^ stromal cells seven days after immunization (Fig 3a, b) and a 6-fold reduction in the absolute number of MRCs at the same timepoint (Fig 3b). To confirm these findings were consistent across mouse strains, we also immunized C57BL/6 mice and determined the MRC response at day seven (Fig 3d-f). We found a significant reduction in both the proportion (Fig 3d, e) and absolute number (Fig 3f) of MRCs, which directly correlated with the poor GC and FDC responses (Fig 1).

**Figure 3.**
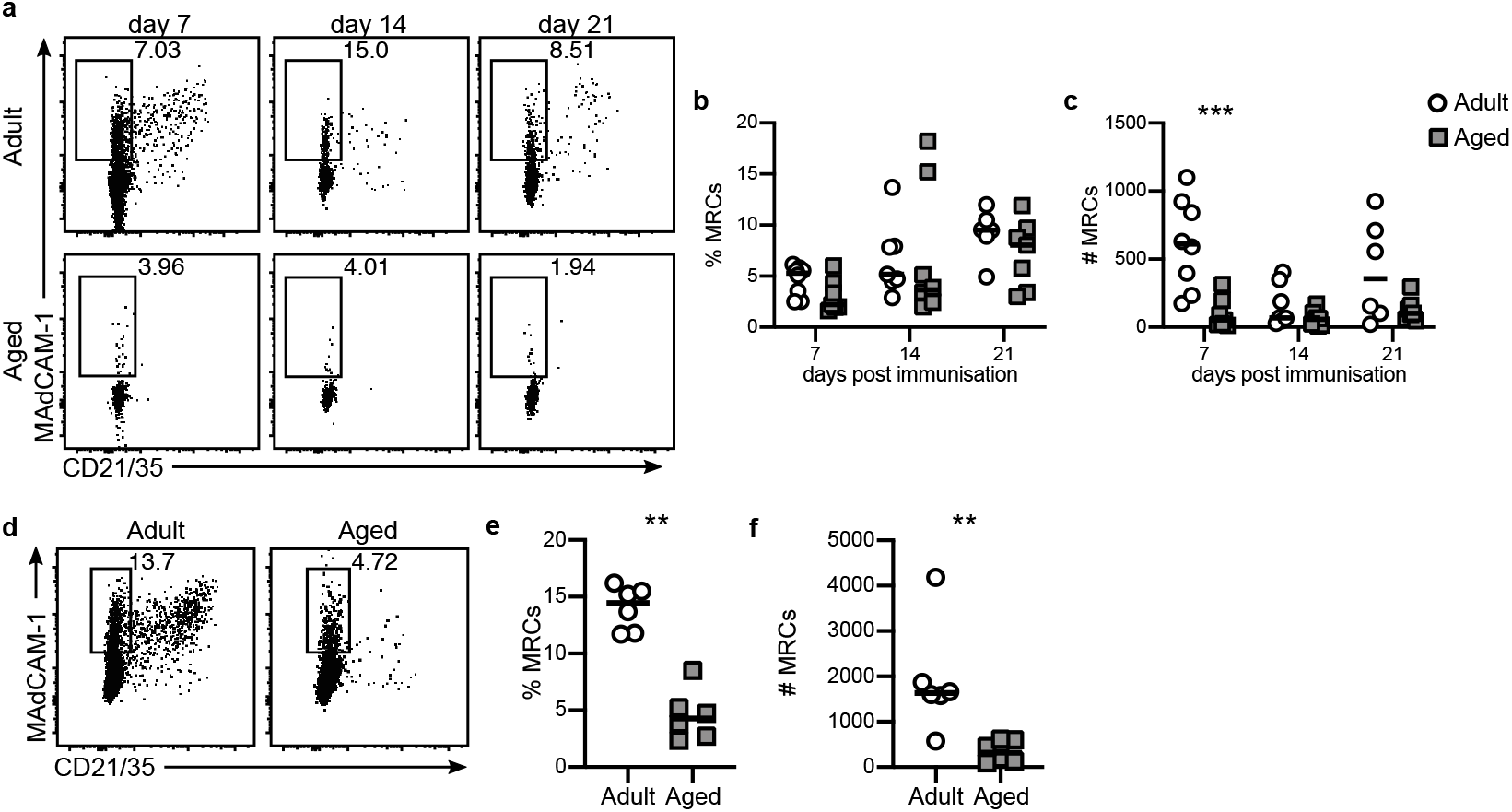
Aging diminishes the MRC response to immunization. Adult (8-12 week old) and aged (>98 week old) mice were immunized subcutaneously with 50μg NP-KLH emulsified in Alum and the MRC response was determined. (a-c) MRC responses were determined in Balb/c mice seven, 14 and 21 days after immunization. (a) Representative gating of MRCs (MAdCAM-1^+^CD21/35^-^) amongst CD31^-^ CD45^-^Pdpn^+^ stromal cells. (b-c) The proportion (b) and absolute number (c) of MRCs. (d-f) MRC responses were determined in C57BL/6 mice seven after immunization. (d) MRCs were gated based on MAdCAM-1 staining amongst CD31^-^CD45^-^Pdpn^+^ stromal cells, and the proportion (e) and absolute number (f) of MRCs was determined. Each symbol represents a biological replicate and lines indicate the median. Data are representative of at least two independent experiments with 6-8 mice per group. Statistical significance was determined using a Mann-Whitney rank test. * p<0.05, **p<0.01, ***p<0.001.

These findings led us to hypothesize that aging negatively impacts the capacity of MRCs to become FDCs, ultimately leading to impaired stromal cell expansion that is associated with poor GC formation. In order to test this, we performed bulk RNA sequencing of sort-purified MRCs, isolated from adult and aged lymph nodes both before and after immunization, based on MAdCAM-1 expression and a lack of CD21/35 expression amongst Pdpn^+^CD31^-^CD45^-^ stromal cells. MRC identity was confirmed by the expression of markers typically associated with this cell type^18^, and lack of immune and endothelial cell-related markers (Extended data Fig 1a). MRCs expressed high levels of *Pdpn*, *Madcam1*, *Ccl19* and *Il7* compared to FDCs (Extended data Fig 1d), as expected of MRCs. Adult MRCs obtained after immunization clustered distinctly from the other three groups by the first principle component (Fig 4a), suggesting that the transcriptome of adult responding MRCs is distinct from cells from the naïve and aged mice. Interestingly, when PC1 was plotted against PC3 we could observe separation of the different MRC groups (Fig 4b), demonstrating that both age and immunization affected the transcriptome of MRCs. In the resting state, 122 genes were differentially expressed between aged and adult MRCs, with 25 genes more highly expressed in adult MRCs and 97 genes more highly expressed in aged MRCs (Fig 4c, Extended data Fig 3a, b). In response to immunization, the expression of 923 genes are significantly altered in adult MRCs: 378 are upregulated and 545 genes are downregulated after immunization (Fig 4d, Extended data Fig 3a, b). In comparison, just 101 genes are differentially expressed in aged MRCs (Fig 4e, Extended data Fig 3e, f). As observed for FDCs, very little overlap was observed between adult and aged MRCs in the genes that altered expression upon immunization (Fig 4f), suggesting that age disrupts the transcriptional response of MRCs to immunization. In order to understand this further, we generated a list of genes that were differentially expressed upon immunization in adult MRCs but were not altered after immunization in aged MRCs and performed gene ontology analysis (Fig 4g, h). Amongst gene ontology terms that were downregulated upon immunization, we saw a striking enrichment of genes associated with extracellular matrix synthesis and organization (Fig 4h, cyan), suggesting that MRCs may alter the local ECM to aid tissue remodeling, as has been described for fibroblastic reticular cells^42^. A number of gene ontology terms were significantly enriched in adult, but not aged, MRCs following immunization (Fig 4h). These can largely be classed as ‘immune response’ (magenta) or ‘initiation of proliferation’ (yellow). This suggested that MRCs proliferate in response to immunization, but that in the aged setting MRCs fail to initiate immune stimulatory and proliferative responses, which may impact on the development of a stromal cell network that is able to support a GC response.

**Figure 4.**
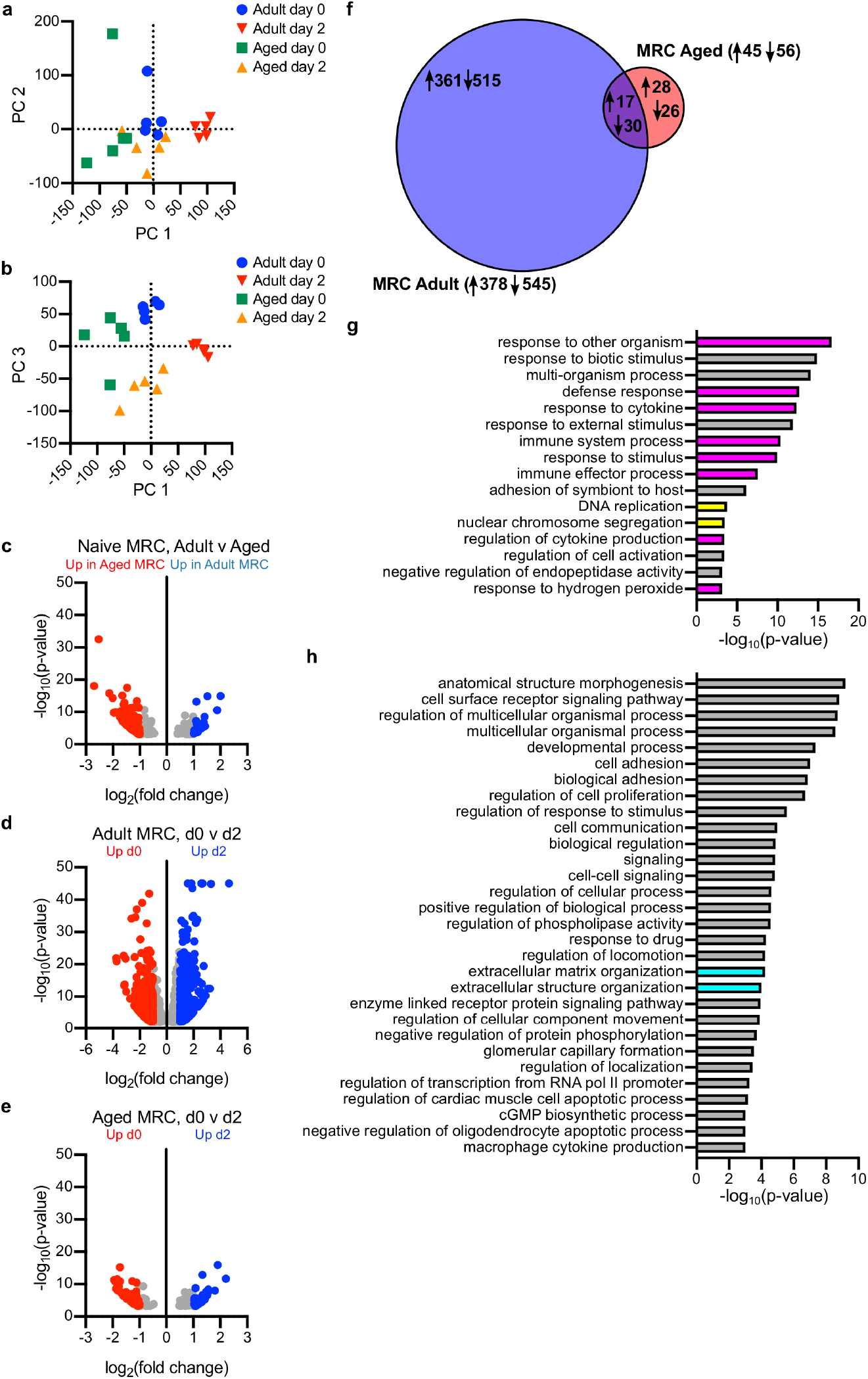
Transcriptional profiling of the MRC response to immunization in aging. MRCs (CD21/35^-^MAdCAM-1^+^Pdpn^+^CD31^-^CD45^-^) were isolated from adult (8-12 week old) and aged (>98 week old) unimmunized mice, or mice immunized two days prior with 50μg NP-KLH emulsified in Alum and subjected to RNA sequencing. (a-b) PCA of all expressed genes, showing PC1 v PC2 (a) and PC1 v PC3 (b). (c) Volcano plots of all genes that are differentially expressed (DESeq2) between adult and aged MRCs isolated from unimmunized mice. Genes that are enriched in aged (red) or adult (blue) are highlighted. (d, e) Volcano plots of all genes that are differentially expressed between naïve and immunized MRCs in adult (d) and aged (e) mice. Genes that are significantly enriched in unimmunized (red) or immunized (blue) MRCs are highlighted. (f) Comparison of differential gene expression following immunization in MRCs isolated from adult or aged mice, where the overlap shows genes that change in both adult and aged MRCs. Numbers indicate genes that are up/down regulated. (g, h) Gene set enrichment analysis was conducted using GOrilla and Revigo to identify gene sets that are uniquely enriched in adult MRCs after immunization. Shown are gene sets that are increased (g) or decreased (h) between d0 and d2 in adult MRCs at d2 but not changed between d0 and d2 in aged MRCs. Five biological replicates were sequenced for each group.

### Failure to initiate MRC proliferation underpins the poor GC response to immunization

To investigate whether aging negatively impacted on the capacity of MRCs to proliferate in response to immunization, we first determined the timecourse of MRC proliferation after immunization in adult mice. We found that MRC proliferation peaked four days after immunization (Fig 5a), in line with that previously described for MRCs^25^, and a time point that is prior to GC formation indicating that MRC remodeling upon immunization precedes GC establishment. We then determined whether MRCs in aged mice could proliferate four days after immunization, based on the peak of the response in adult mice. We found that, as indicated by our sequencing results (Fig 4), a lower proportion of MRCs from aged mice expressed ki67 (Fig 5b, c), indicating fewer proliferating cells. When we enumerated MRCs isolated four days after immunization, we found they were reduced by proportion (Fig 5d, e) and absolute number (Fig 5f) in aged mice. FDCs were also reduced in proportion (Fig 5d, g) and absolute number (Fig 5h) in aged mice; this reduction was so pronounced that FDCs were often not isolated from aged lymph nodes at this timepoint. We also observed a reduction in the activation of MRCs (Fig 5i, j), as measured by the mean fluorescence intensity (MFI) of cell surface Pdpn staining^43^, suggesting these cells were not as efficiently activated in an aged setting.

**Figure 5.**
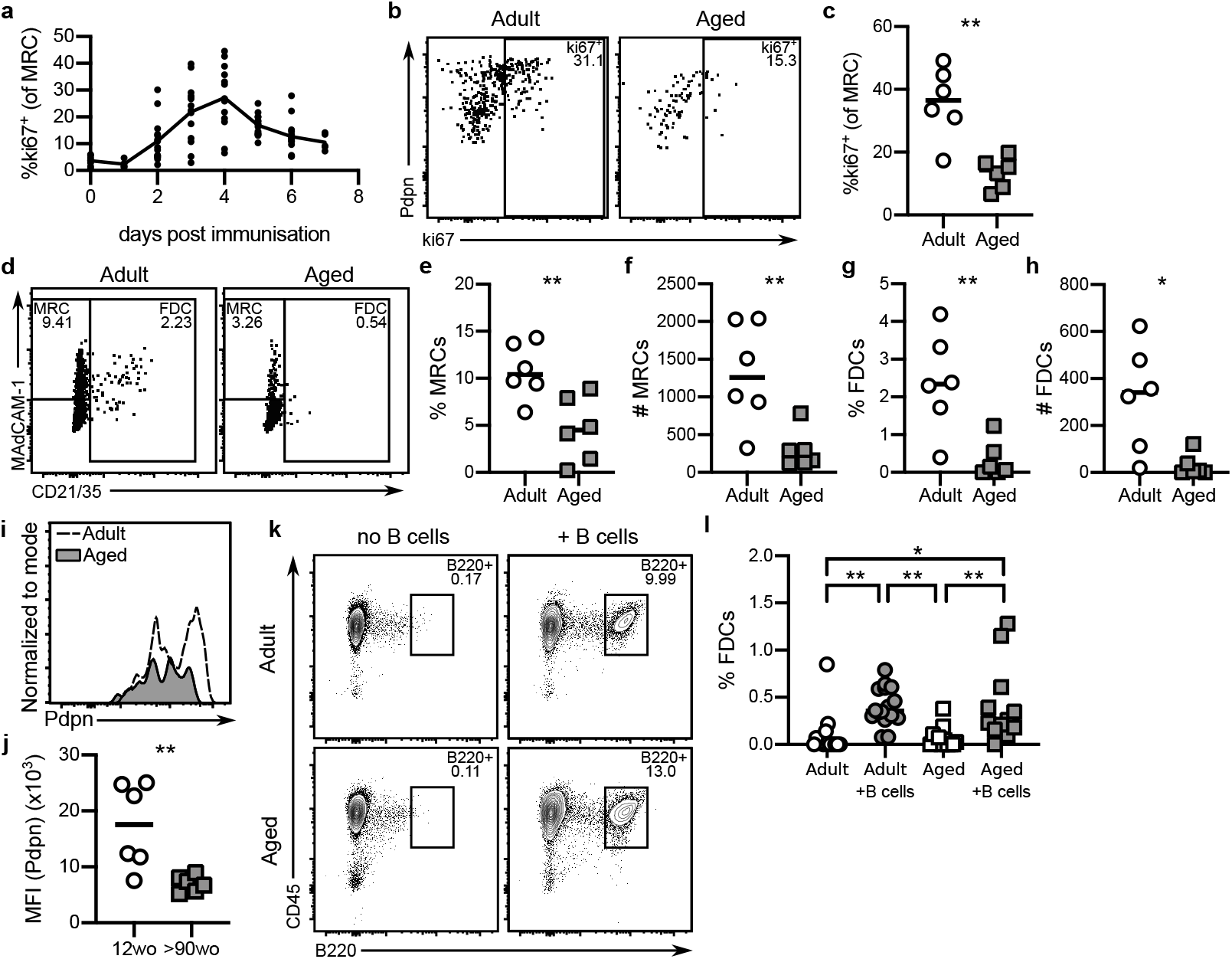
Aged MRCs fail to proliferate in response to immunization. (a) The proliferation of MRCs in response to immunization was determined over a timecourse in adult mice. (b-c) The proliferation of MRCs four days after immunization in adult and aged C57BL/6 mice was determined by ki67 staining (b) and measured as a proportion of MRCs (c). (d-h) The expansion of MRCs and FDCs was determined four days after immunization in adult and aged mice by MAdCAM-1 and CD21/35 staining (d). Both the proportion (e, g) and number (f, h) of MRCs (e, f) and FDCs (g, h) were determined. (i-j) The activation of MRCs was also measured by flow cytometry (i) and determined by the mean fluorescence intensity of Pdpn (j). (k-l) Purified C57BL/6 B cells isolated from adult mice were adoptively transferred into adult (8-12 week old) and aged (>95 week old) B cell-deficient (μMT^-/-^) mice (k). Seven days later the development of FDCs (CD21/35^+^) was determined as a proportion of Pdpn^+^CD31^-^CD45^-^ stromal cells (l). Data in (b-j) are representative of three independent experiments with 5-8 mice per group. Statistical significance was conducted using a Mann-Whitney rank test. * p<0.05, **p<0.01. Data in (a) are combined of three separate experiments comprising 4-6 mice per timepoint. Data in (k, l) are combined of three independent experiments, comprising 3-5 mice per group. Statistical significance was determined using a one-way ANOVA with Tukey’s multiple comparison test. Each symbol represents a biological replicate and lines indicate the mean. * p<0.05, **p<0.01.

Because FDCs can be derived from MRCs^25^, we next sought to determine whether aged MRCs have the capacity to differentiate into FDCs in general or whether this was a defect specific to vaccination. FDC development in lymph nodes requires lymphotoxin-beta receptor signaling from B cells^44, 45^ and B cell-deficient μMT^-/-^ mice lack FDCs in adult lymph nodes. Adoptive transfer of B cells leads to FDC differentiation, and this is independent of MRC proliferation^25^. We used the same model to understand MRC-to-FDC differentiation in aging by adoptively transferring B cells purified from wildtype adult mice into adult (7 week old) or aged (>95 week old) μMT^-/-^ mice. There was no difference in the engraftment of B cells between adult and aged lymph nodes (Fig 5k), and the development of FDCs, determined seven days after transfer, was similar between adult and aged mice (Fig 5l), suggesting that the lymphotoxin-beta receptor-dependent capacity of MRCs to differentiate into FDC was not affected by age. Taken together these data suggest that the activation and proliferation of MRCs induced by immunization is uniquely diminished in an aged setting, resulting in fewer FDCs within the GC after immunization.

### The size of the FDC network is a key contributor to GC output

In the absence of an *in vivo* tool to specifically reduce the FDC network in mice, we used the *hyphasma* mathematical modeling approach^46, 47^ to understand whether a reduced FDC network would be predicted to alter GC size or output. In order to accurately model how the age-associated changes in the size of the FDC network impact the GC response, we first determined the size of the FDC network *in vivo*: immunostaining on cryosections of lymph nodes obtained 14 days after immunization of adult (8-12 week old) and aged (90-95 week old) Balb/c mice (Fig 6a) and determined the spatial changes that occur to the GC with age. Both the FDC area (Fig 6b), as measured by the cross-sectional CD35^+^ area within IgD^-^ GCs, and the proportion of the GC that was occupied by FDCs (Fig 6c) were significantly reduced (1.7-fold) in the lymph nodes from aged mice.

**Figure 6.**
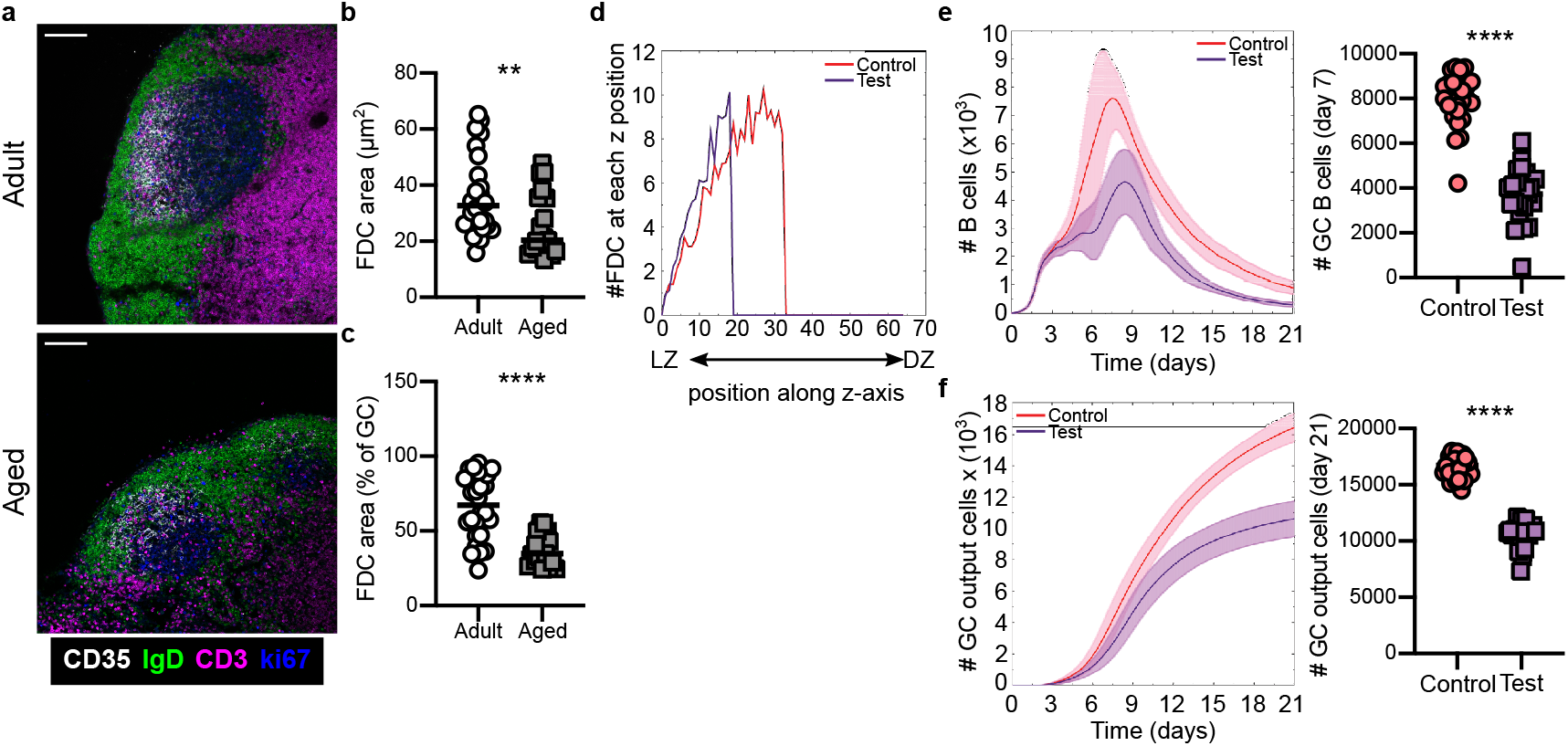
Mathematical modeling of the FDC network size impacts on GC output. (a-c) Cryosections from lymph nodes obtained 14 days after immunization of 12 week old (adult) or >90 week old (aged) Balb/c mice were stained for CD35 (white), IgD (green), CD3 (magenta) and ki67 (blue). Shown are representative images (a) and calculations of the cross-sectional area of the FDC network (b) and the proportion of the GC that is occupied by FDCs (c). Each dot represents a single GC, with data compiled from five individual mice per group. Statistical significance was conducted using a Mann-Whitney rank test. **p<0.01, ****p<0.0001. Scale bars 100μm. (d-f) Mathematical modeling of the impact of reduced FDC area on the generation of GC B cells using *hyphasma*. (d) Spatial representation of the number of FDCs and positioning along the z-axis. (e) Total number of B cells generated through the GC reaction. Left: model output, right: graphical representation of number of GC B cells at day seven. (f) Total number of output GC B cells. Left: model output, right: graphical representation of number of output GC B cells at day 21. Lines in kinetic plots represent the mean of 25 simulations and shaded areas indicate the standard deviation. For graphs, each dot represents a simulation and lines indicate the median. Statistical significance was conducted using a Kruskal-Wallis test with Dunn’s multiple testing correction. ****p<0.0001.

To model the reduction of FDC in the aged setting we reduced the size of the FDC network by decreasing both the proportion of the GC that was covered by FDCs (control = 50%, test = 30%) and the number of FDCs (control = 200 cells, test = 100 cells). We did not alter the total level of antigen on the surface of FDCs, allowing us to determine the effect of FDC network size independent of antigen capture. These changes maintain the same FDC density, resulting in FDCs being positioned closer to the light zone (Fig 6d), thus mimicking the reduction in GC light zone (Fig 6a-c) and the reduction in the number of FDCs (Fig 1) observed in aged mice. Reducing the FDC network has a significant impact on the GC: the number of GC B cells at day seven (Fig 6e) and the number of GC output cells at day 21 (Fig 6f) are reduced when the FDC network resembles the aged state. This phenocopies our experimental data from aged mice (Fig 1), suggesting that poor expansion of the FDC network contributes to the impaired GC response in aged mice independently of antigen capture and retention.

### The poor aged GC response is primarily driven by defects in the aged lymph node microenvironment

The data above demonstrate that MRCs are poorly responsive to vaccination in aged mice, and this is linked to poor expansion of the FDC network. Mathematical modeling implicates these stromal cell defects as causal in the defective GC in ageing. In order to test this prediction *in vivo* we established a heterochronic parabiosis model in which the effect of age on lymphocytes can could be distinguished from the effect of age on the lymph node stroma (Fig 7a). Parabiosis is a procedure through which the circulatory system of animals is surgically conjoined, generating a system in which blood-derived cells are shared across individual mice and non-migratory cells, such as stromal cells, remain that of the original host^48^. Thus, this is an ideal system through which we can interrogate the impact of the aged microenvironment on GC formation.

**Figure 7.**
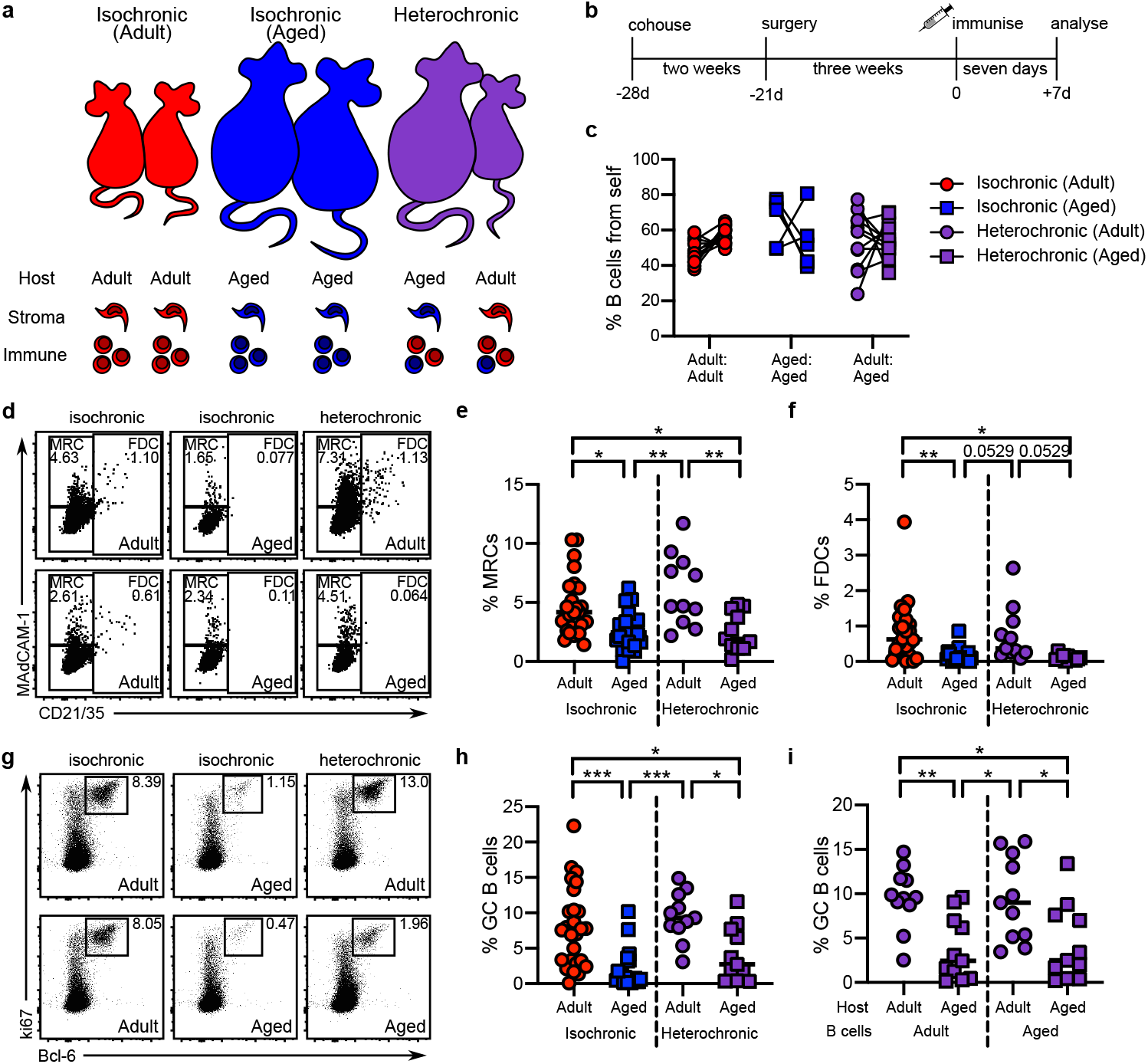
The poor GC and stromal cell response in aging is driven by the lymph node microenvironment. Heterochronic parabiosis was used to determine the role of the aged lymph node microenvironment in the generation of the GC response. Congenically distinct (CD45.1^+^ or CD45.2^+^) C57BL/6 mice were paired in isochronic (adult:adult, red; or aged:aged blue) parabionts or heterochronic (adult:aged, purple) parabionts. Adult mice (7-12 weeks old) are represented with circles; aged mice (>92 weeks old) are represented with squares. (a) Depicts the experimental groups and the age of relevant cell types in the lymph node in each group. (b) Experimental protocol: mouse pairs were cohoused for two weeks, then surgically joined; three weeks after surgery mice were immunized with 50μg NP-KLH emulsified in Alum subcutaneously on the outer flanks; seven days after immunization the stromal and GC responses were determined by flow cytometry. (c) Successful parabiosis was determined by the blood B cell chimerism; mice with 40-80% self-derived B cells were used for further analyses. (d-f) The stromal cell responses were measured using MAdCAM-1 and CD21/35 staining (d); the proportion of MRCs (MAdCAM-1^+^CD21/35^-^) (e) and FDCs (CD21/35^+^) (f) were determined as a proportion of CD31^-^CD45^-^Pdpn^+^ stromal cells. (g-i) The GC response was measured by ki67^+^Bcl-6^+^ staining (g), and determined as a proportion of B220^+^ B cells (h). Within heterochronic parabionts, B cells were gated based on age using congenic markers, and their capacity to generate a GC response within adult or aged hosts was determined (i). Data are combined from six separate experiments, using 4-20 mice per group (2-10 pairs). Data in (c) show individual mice with lines connecting paired blood samples. In (e, f, h, i) each data point represents an individual mouse; isochronic parabionts are represented as a single group. Responses to immunization are independent; blood samples are paired. Statistical significance was determined using a two-way ANOVA with Sidak’s multiple comparison test (c) or a one-way ANOVA with Holm-Sidak’s multiple comparison test (e, f, h, i). * p<0.05, **p<0.01, ***p<0.001.

We generated three groups of parabiotic pairs using congenically distinct mice: an adult isochronic group, where adult mice were parabiosed with mice of the same age (6-12 weeks; red); an old isochronic group, where aged mice were parabiosed with aged mice (>92 weeks; blue); and a heterochronic group, where younger adult (6-12 weeks) mice were parabiosed with aged (>92 weeks) mice (purple) (Fig 7a). Mice were cohoused in pairs for two weeks prior to surgery to allow for social conditioning and rested for three weeks after surgery to allow revascularization and establishment of parabiosis. Mice were immunized on outer flanks (contralateral to the surgical site) and seven days after immunization we determined stromal cell responses and GC formation (Fig 7b). Mice become T cell lymphopenic with age^49^, meaning that in heterochronic pairs, adult-derived T cells typically predominate in both mice, while aged:aged mice are often mismatched for T cell parabiosis due to variability in T cell lymphopenia (Extended Data Fig 4). Therefore, the degree of parabiosis was determined by CD45.1:CD45.2 chimerism of B cells in the blood of parabiotic pairs (Fig 7c). We found no significant difference in the establishment of blood parabiosis between the groups, although it is important to note that the aged isochronic group was variable.

We first sought to determine whether the stromal cell response to immunization was altered by the presence of lymphocytes from an animal of a different age by measuring the expansion of MRCs and FDCs as a proportion of CD45^-^CD31^-^Pdpn^+^ stromal cells (Fig 7d) in adult:adult isochronic, aged:aged isochronic and adult:aged heterochronic mice. The expansion of MRCs (Fig 6e) and FDCs (Fig 6f) was significantly diminished in the aged isochronic mice, compared to the adult isochronic mice, in line with our previous findings in aged mice (Fig 1, Fig 3). In heterochronic parabionts, we observed that the expansion of MRCs was significantly diminished the aged mouse compared to its adult partner (Fig 7e). A similar trend was observed for the expansion of FDCs in heterochronic parabionts, where FDCs fail to expand in aged mice (Fig 7f). Thus, the age of the lymph node stromal cells diminishes their ability to respond to immunization, and this cannot be rescued by exposure to lymphocytes derived from younger adult mice.

We then sought to determine whether the age of stromal cells contributes to the magnitude of the GC. We measured the size of the total GC response in adult:adult isochronic, aged:aged isochronic and adult:aged heterochronic mice by determining the proportion of ki67^+^Bcl-6^+^ GC B cells amongst B220^+^CD4^-^ B cells (Fig 7g). Consistent with the observed age-associated defect in GC formation (Fig 1), the aged isochronic pairs had fewer GC B cells than adult isochronic pairs (Fig 7h). In heterochronic pairs, the magnitude of the GC reaction was dictated by the age of the host: the GC response was significantly reduced in aged animals, despite the B cell pool deriving from both adult and aged mice (Fig 7h). This suggests that the age of the microenvironment dictates the size of the GC response. Because the mice are congenically marked (CD45.1^+^ or CD45.2^+^), we can use the heterochronic parabionts to determine whether B cell-intrinsic aging influences their capacity to participate in the GC by measuring the ability of immune cells from younger adult mice to generate a GC response in both adult and aged lymph nodes. We found that B cells from adult mice were impaired in their capacity to generate GC B cells in an aged lymph node (Fig 7i). In agreement with this, B cells from aged mice were as able to participate in the GC response as B cells from younger adult mice if they were in a younger lymph node (Fig 7i). These data suggest that the age of the lymph node microenvironment, rather than that of the B cells, dictates their capacity to generate GC responses in an aged setting. Together with the *in silico* modeling data (Fig 6), this demonstrates that aging causes a cell-intrinsic defect in lymph node stromal cells that is the primary contributor to the impaired GC response observed in older individuals.

## Discussion

The diminution of the GC response with advancing age is associated with poor protective immunity in infections and vaccine settings^2, 3, 4^. Understanding the fundamental age-associated mechanisms that drive this response are critical to inform vaccine design to induce optimal protective immunity in older individuals. We show here that GC and FDC responses to immunization are impaired in aged mice, and that this is driven primarily by the poor cellular and molecular response of MRCs to immunization. Using in silico modeling we demonstrate that a reduction in the number of FDCs contributes to the poor GC response. Finally, using heterochronic parabiosis we show that the age of the lymph node microenvironment is a primary driver of poor vaccine-induced GC responses in aging. Together our data uncover a new paradigm in aging immunity: poor responsiveness of MRCs to immunization disrupts the expansion of GC-associated stromal cell networks, underpinning the impaired GC responses observed with advancing age.

The human lymph node becomes increasingly disrupted with advancing age, and the GC response is the most affected compartment, where both the number of GCs and the size of each GC is reduced^50, 51, 52, 53^. B cell age does not largely alter their capacity to participate in a GC^33^, and our data confirm these findings. In contrast to B cells, T cell responses are impaired in older individuals. Notably, the magnitude of the Tfh cell response is reduced in mice and humans^34, 54^, and their ability to support GC formation^30^ and B cell selection^33^ is impaired by age. T cell age is, however, not the only contributor to impaired GC responses – transfer of young T cells into aged T cell-deficient mice results in a diminished GC response^36^, consistent with a role for the defective stromal cell response underpinning poor GCs in ageing. Importantly, age-associated degeneration of lymph node structure has functional consequences, and is intrinsic to the microenvironment: poor T cell immigration and survival is observed in aged lymph nodes, and this is not rejuvenated by circulatory factors in heterochronic parabiosis^55^.

FDCs are central to GC structure and function^8^, and studies have shown that FDC networks poorly expand in aged mice^56^. Aged FDCs are also less able to support B cell responses *in vitro*, even when cultured with T and B cells derived from younger adult mice^57^. Studies of FDCs in aging have suggested that their inability to support GC formation is driven by reduced antigen capture and retention^56^, which may be a consequence of reduced Fc receptor expression^40^. Our data extends these observations, demonstrating that age limits the expansion of FDCs, and adds the conceptual advance that the MRC response is the primary driver of these defects. Age impairs the MRC response, which is characterized by upregulation of genes associated with immune activation, such as chemokines and cytokines, as well as proliferation, and results in poor FDC differentiation.

MRCs are a poorly defined lymph node stromal cell subset. They reside beneath the subcapsular sinus, overlaying the B cell follicle and the interfollicular regions and form a reticular layer that retains stem-like properties^23, 24^. It has been postulated that MRCs play a role in antigen transport to the follicle, based on their location beneath the subcapsular sinus. Studies of antigen drainage have shown that, in aging, antigen fails to access the follicle and remains trapped beneath the subcapsular sinus^58^, held by reticular-like cells that are morphologically similar to FDCs, and form a connected chain between the subcapsular sinus and the FDC network in the follicle^59^, and are likely synonymous with MRCs. Taking these studies together with our data we can build a model of how stromal cell dysfunction impedes GC development in aging: MRCs are poorly activated in response to immunization, impeding their proliferation and differentiation into FDCs and resulting in antigen being retained – or trapped – on MRCs adjacent to the subcapsular sinus. A mechanism through which MRCs may capture antigen has not been described, but may relate to their upregulation of complement receptors during differentiation into FDCs^25^. Ultimately, reduced FDCs differentiation, poor antigen access to the follicle, and limited antigen capture by FDCs together impede GC development independently of an effect of immune cell-intrinsic age.

MRCs also play a role in regulating the subcapsular sinus niche^60^ and are thought to support the localization and survival of innate lymphoid cells in the interfollicular niche through provision of interleukin-7^61^. The interfollicular region is also an important site for the initiation of humoral immunity^62^ in part through the recruitment of NKT cells that promote humoral immunity^63^, and our transcriptomic data suggest that MRC activation may play a role in these phenomena, for example through upregulation of chemoattractant or immune stimulating molecules such as *Ccl20* and *Il33*. MRC-like stromal cells in the interfollicular region are also key to anti-helminth responses, promoting *de novo* B cell follicle formation^64^ and potentially contributing to T cell-mediated type 2 immunity^65^. Intriguingly these MRC-like cells do not differentiate into FDCs in the newly developed follicles^64^, suggesting that FDC differentiation is not always the ultimate fate of stimulated MRCs. FDCs are dependent on lymphotoxin beta receptor signaling for development and survival^44, 45^, and lymphotoxin signaling has been shown to drive FDC activation and GC formation during immunization^66^. This pathway is, however, unlikely to play a role in aging: we do not find an age- or immunization-associated change in the level of *Ltbr* expressed by stromal cells, and the exposure of aged stromal cells to younger adult T and B cells does not boost their response to immunization. This suggests that other factors drive the activation of MRCs and their subsequent differentiation into FDCs. How MRCs respond to and integrate immune and pathogen-driven signals is not known, and it is tempting to speculate that lymph node stromal cells can act as a rheostat for the induction of adaptive immune responses by directing the development of niches that support pathogen-specific immunity.

In summary, our study demonstrates that the MRC-to-FDC differentiation pathway is a critical step in GC formation, and that disruption of this program in aging impedes GC formation in a manner that represses humoral immunity. Targeting stromal cells offers a novel avenue through which vaccine efficacy could be improved in older individuals. Further studies into lymph node stromal cell behavior will discover pathways that may be exploited to manipulate GC responses for therapeutic benefit.

## Material and Methods

### Mice

C57BL/6, Balb/c, μMT^-/-^ and *Fap*^tT^A::*TetO*^cre^::*Rosa26*^lox-stop-lox-tdTomato 41^ mice were bred and housed in the Babraham Institute Biological Support Unit. No primary pathogens or additional agents listed in the FELASA recommendations were detected during health monitoring surveys of the stock holding rooms. Ambient temperature was 19-21°C and relative humidity 52%. Lighting was provided on a 12-hour light: 12-hour dark cycle including 15 min ‘dawn’ and ‘dusk’ periods of subdued lighting. After weaning, mice were transferred to individually ventilated cages (GM 500: Techniplast) with 1-5 mice per cage. Mice were fed CRM (P) VP diet (Special Diet Services, #801722) ad libitum and received seeds (e.g., sunflower, millet) at the time of cage-cleaning as part of their environmental enrichment. All UK mouse experimentation was approved by the Babraham Institute Animal Welfare and Ethical Review Body and the UK Home Office. Animal husbandry and experimentation was conducted in accordance with existing European Union and United Kingdom Home Office legislation and local standards. Parabiosis experiments were conducted at the Animal Research Centre, University of Leuven, Belgium using aged C57BL/6 mice purchased from Charles River (UK) and C57BL/6 and CD45.1^+^ mice bred and aged in-house. All Belgian mice were housed under specific pathogen-free conditions and fed ssniff® R/M-H chow (4.7% w/v sugar, 3.3% w/v fat, 19% w/v protein) and were used in accordance the University of Leuven Animal Ethics Committee and existing European Union standards. Adult mice were 8-12 weeks old and aged mice were 90-105 weeks old when used for experiments. All experiments were conducted use sex-matched controls, and both male and female cohorts were used.

### Immunizations

Mice were immunized with 50μg of NP-KLH (4-hydroxy-3-nitrophenylacetyl-Keyhole Limpet Hemocyaninin; conjugation ratio 29-33, Biosearch technologies N-5060) dissolved in PBS at 2mg/mL and emulsified at a 1:1 ratio with Imject Alum Adjuvant (ThermoFisher Scientific 77161). Mice received 50μL subcutaneously on each flank at the inguinal lymph node site under inhalation anesthesia. Lymph nodes were harvested after two, four, seven, 14 or 21 days and processed for analysis. Parabionts were immunized on the upper and lower flanks, draining the axillary and brachial lymph nodes as well as the inguinal site. Lymph nodes in parabiosis experiments were harvested seven days after immunization. Blood samples were collected by cardiac puncture at the time of dissection to determine blood chimerism for parabiotic pairs.

### Parabiosis

Parabiosis was conducted essentially as described^48, 67, 68^. Congenically distinct (CD45.1 and CD45.2) female C57BL/6 mice were cohoused for two weeks prior to surgery to ensure harmonious cohabitation. Surgery was conducted under inhalation anesthesia according to LASA Dhiel guidelines. One incision was made from elbow to knee on corresponding lateral sides and the skin separated from the subcutaneous fascia by blunt dissection. The right olecranon of one mouse was attached to the left olecranon of the second mouse with a single suture; this was repeated for the knee joint. The dorsal and ventral skins were anastomosed by continuous suture. Mice we administered analgesia (10mg/kg Carprofen) intraperitoneally twice daily for 48hr. Parabionts were housed as single pairs with free access to food and water and monitored daily for clinical signs. Mice that appeared in distress (hunching, piloerection) were humanely killed. Three weeks after surgery, mice were immunized as above on the outer flanks. Seven days after immunization, blood and lymph node samples were collected and analyzed. Mice that failed to parabiose (<20% of B cells from the partner mouse) were excluded from further analysis.

### Cell isolation

For analysis of stromal cells, single cell suspensions were generated by enzymatic digestion as described^13, 41^. Briefly, lymph node capsules were pierced with fine forceps and incubated with 0.5mL digestion buffer at 37°C for 20mins and tubes inverted every 5min. After 20mins lymph nodes were triturated with a 1mL pipette tip and supernatant containing released cells was collected to ice-cold buffer comprising PBS containing 2% FCS and 2mM EDTA. Remaining lymph node fragments were digested twice more as described, triturating with a 1mL pipette at 5min intervals. Digestion buffer comprised 0.2mg/mL Collagenase P (Sigma, #11213865001), 0.8 mg/mL Dispase II (Sigma #4942078001) and 0.1mg/mL DNase I (Sigma, #10104159001) in RPMI 1640 media (Gibco, #11875093). Cell suspensions were passed through a 70μm mesh prior to staining. For analysis of GC B cells, lymph nodes were forced through a 40μm mesh in PBS containing 2% FCS. In some experiments GC B cells were analyzed from single cell suspensions prepared by enzymatic digestion, as for stromal cell analysis.

### Flow cytometry

Cells were stained using cocktails of antibodies. Fc receptors were blocked prior to staining using anti-CD16/32 (93, ThermoFisher Scientific). Where >1 brilliant violet or brilliant UV dye was used concurrently, cells were stained using BD brilliant stain buffer (BD Biosciences #563794). Cells were fixed and permeabilized for intracellular/nuclear staining using the eBioscience Foxp3/ Transcription Factor Fixation/Permeabilization Staining buffer set (ThermoFisher Scientific, #00-5523-00). Samples were stained for 1hr at 4°C (extracellular) or room temperature (intracellular/nuclear) and fixed/permeabilized for 30mins at 4°C. Antibodies used were: CD45 (30-F11, BioLegend), CD31 (MEC13.3, Biolegend or BD Biosciences), Pdpn (8.8.1, ThermoFisher Scientific), MAdCAM-1 (MECA-367, BioLegend), CD21/35 (7E6, BioLegend; eBio8D9, ThermoFisher Scientific), ICAM (YN1/1.7.4, BioLegend), VCAM (MVCAM.A, BioLegend), CD44 (IM7, BioLegend), ki67 (SolA15, ThermoFisher Scientific), B220 (RA3-6B2, BioLegend), CD4 (GK1.5 or RM4-5, BioLegend or ThermoFisher Scientific), Bcl-6 (K112-91, BD Biosciences). In some experiments biotinylated antibodies were detected with BUV-395-conjugated streptavidin (BD Biosciences, #564176). Dead cells were excluded with Fixable live/dead e780 (ThermoFisher Scientific, #65-0865-18). Samples were acquired on a BD LSRFortessa (BD Biosciences) harbouring 355, 405, 488, 561 and 640nm lasers and analyzed using FlowJo software (TreeStar).

### Immunofluorescence and confocal microscopy

Lymph nodes were immersion-fixed in periodate-lysine-paraformaldehyde containing 1% PFA (Sigma #P6148), 0.075 M L-lysine (Sigma #L5501), 0.37 M sodium phosphate (pH 7.4, Sigma #342483), and 0.01 M NaIO4 (Sigma #210048) for five hours at 4°C, washed, cryoprotected in 30% sucrose (Sigma #S0389) overnight at 4°C, embedded in optimal cutting temperature medium (VWR #25608-930) on a dry ice/ethanol bath and stored at −80°C. Sections (20μm) were cut at −16°C on a cryostat (Leica), air-dried overnight and stored at −20°C (short-term) or −80°C (long-term). For staining, sections were defrosted and air-dried overnight at room temperature, rehydrated with PBS containing 0.05% Tween-20 (PBST), permeabilised with 2% Triton X-100 (Sigma #X100) for 15min at room temperature and blocked in PBST containing 1% BSA for 1hr at room temperature. Tissues sections were stained with antibodies directed against CD35-biotin (8C12, BD Biosciences), ki67-FITC (SolA15, ThermoFisher Scientific), IgD-AF594 (11-26c.2a, BioLegend) and CD3-APC (17A2, ThermoFisher Scientific) overnight at 4°C. Biotinylated CD35 was detected with BV421-conjugated streptavidin (BioLegend, #405225) for 2hr at room temperature. Sections were washed, mounted in aqueous mounting media (HydroMount; National Diagnostics, #HS-106) and allowed to dry prior to imaging. Sections were obtained using a Zeiss LSM880 harboring 405-, 432-, 488-, 561-, and 633-nm lasers using 20×/0.50 air or 40×/1.40 oil objectives. Image channels were collected separately and analyzed and compiled using Fiji (National Institutes of Health) software. FDC and GC areas were calculated using CellProfiler^69^: the GC area (μm^2^) was defined as IgD^-^ki67^+^CD35^+^; analysis was performed using automated identification of GCs, with the light zone FDC network area (μm^2^) defined as IgD^-^CD35^+^.

### Mathematical modeling of GC output

*In silico* mathematical modeling was carried out using the *hyphasma* model, as described in^46^. Briefly, the GC is modeled as a grid within a sphere; each grid point is the size of a cell (5μm) and has a defined concentration of the dark zone and light zone chemokines, CXCL12 and CXCL13, respectively. Each simulated cell’s properties, that is position, polarity, velocity, antigen receptor, and amount of internalized antigen, are updated at each time-step and can be monitored computationally. The direction of movement is described by a persistent time and a distribution of turning angles that is influenced by chemotaxis. GC B cells are separated into proliferating centroblasts in the dark zone and centrocytes undergo selection in the light zone. The proliferation, death and selection of GC B cells are incorporated as probabilistic events depending on the affinity of the receptor and the interaction with FDCs or Tfh cells. Stellate FDCs are spread randomly in a fraction of the sphere defined as the light zone and the total amount of antigen is spread equally between all grid positions occupied by FDCs. In each simulation, B cells undergo selection based on antigen abundance, antigen uptake and the probability of Tfh cell encounter. Each simulation models the typical GC steps where B cells first search for, and acquire, antigen from FDCs, then search for, and interact with, Tfh cells to receive a proliferation signal before exiting the GC or recycling to the DZ. Tfh cell help is given to interacting B cells with the highest amount of antigen, which also indicates B cell receptor affinity, and the model assumes that centrocytes interact with Tfh cells once per cycle. In the model used here, the GC is seeded with two B cell clones per hour over the first three days, each of which divides six times^70^; this models the *in vivo* findings that 10-100 B cell clones seed a founder GC^71^. A full list of parameters is available in^72^.

### Adoptive transfer

Spleens from 8 week old wildtype mice were forced through a 40μm sieve and B cells were enriched using the MagniSort B cell enrichment kit (ThermoFisher Scientific, 8804-6827-74) according to manufacturer’s instructions. Mice were administered 2-4 million B cells intravenously via the tail vein.

### RNA Sequencing and analysis

Stromal cells were released from lymph nodes by enzymatic digestion as described above. Hematopoietic cells were depleted using anti-CD45 microbeads (Miltenyi Biotec, 130-052-301) (7μL/10^7^ cells), a QuadroMACS magnet (Miltenyi Biotec) and LS columns (Miltenyi Biotec, 130-042-401), according to manufacturer’s instructions. The CD45^-^ fraction was retained and stained with live-dead e780 and antibodies directed against CD45, Pdpn, CD31, MAdCAM-1 and CD21/35 as described above. MRCs (MAdCAM-1^+^CD21/35^-^Pdpn^+^CD31^-^CD45^-^) or FDCs (MAdCAM-1^+/-^CD21/35^+^Pdpn^+^CD31^-^CD45^-^) from unimmunized or immunized adult (8 weeks old) and aged (>95weeks old) mice were sorted directly into 8.5μL of lysis buffer supplied with the SMART-Seq v4 Ultra low Input RNA kit for Sequencing (Clontech #634890). Where feasible we matched the number of sorted FDCs (28-342 in unimmunized, 18-82 in immunized) and MRCs (263-798 in unimmunized, 188-520 in immunized) between adult and aged samples at each time point to limit input cell number variation in the sequencing data.

cDNA from sorted cells was generated using the SMART-Seq v4 Ultra low input RNA kit (Clontech # 634890) according to manufacturer’s instructions. Libraries were prepared with 400pg of cDNA per sample using the Illumina Nextera XT kit (#FC-131-1096) following manufacturer’s instructions. cDNA and sequencing library quality were determined using Agilent BioAnalyser High Sensitivity DNA chips (#5067-4626), Qubit dsDNA High sensitivity Assay kit (Invitrogen #Q32854) on a Qubit 4 fluorometer (Invitrogen), and a KAPA library quantification kit (Roche #7960140001). Samples were sequenced on a HiSeq 2500 (Illumina, v4 chemistry) as 100bp single-end reads. Analysis was performed using the SeqMonk software package (Babraham Institute, https://www.bioinformatics.babraham.ac.uk/projects/seqmonk/, version 1.45.4) after trimming (Trim Galore v0.4.2) and alignment of reads to the reference mouse genome GRCm38 using HISAT2^73^. Reads were quantitated over exons and library size was standardized to reads per million, and then read counts were log_2_ transformed. Data were analyzed by cell type. Expressed genes were defined as those with a log_2_(reads per million)>-4 in at least five biological replicates. Principal component analysis was conducted using all expressed genes. Differentially expressed genes (p<0.05, log2fc>1) were identified using Seqmonk’s built-in DESeq2 analysis^74^ of raw counts in pairwise comparisons: cells from unimmunized adult mice were compared to cells from unimmunized aged mice; and immunized cells were compared to unimmunized cells for both aged and adult mice.

Differentially expressed genes (p<0.05) with a log_2_fc>1 in adult, but not aged, samples were used for gene ontology analysis, which was conducted using Gorilla (http://cblgorilla.cs.technion.ac.il/) against a list of background genes; gene ontology terms were refined using Revigo (http://revigo.irb.hr/), with a small gene list (allowed similarity = 0.5) output.

### Statistics

Experiments were repeated a minimum of two times; in some analyses independent experiments were combined. Data were analyzed and visualized using Prism 8 (GraphPad). Statistical significance was determined using the Mann-Whitney rank test where two groups were compared, or a one-ANOVA with Tukey’s or Holm-Sidak’s multiple testing correction or two-way ANOVA with Sidak’s multiple testing correction where >2 groups were compared. P-values <0.05 were considered statistically significant. In graphs, each data point represents a biological replicate and lines indicate the median.

## Acknowledgements

We thank staff from the Biological Support Unit and Flow Cytometry and Imaging Facilities (Babraham Research Campus) for support.

AED was supported by a Biotechnology and Biological Sciences Research Council Future Leader Fellowship (BB/N011740/1), ASC was supported by European Union’s Horizon 2020 research and innovation programme “ENLIGHT-TEN” under the Marie Sklodowska-Curie grant agreement No.: 675395, DLH was supported by a National Health and Medical Research Council Australia Early-Career Fellowship (APP1139911). This work was supported by an H2020 European Research Council grant to MAL (637801 TWILIGHT), a Biotechnology and Biological Sciences Research Council grants to MAL (BBS/E/B/000C0408 and BBS/E/B/000C0407), a European Research Council Consolidator grant to AL (TissueTreg), a Human Frontier Science Program grant to MMH (RGP0033/2015), and a Biotechnology and Biological Sciences Research Council Campus Capability grant to the Babraham Institute.

## Author Contributions

AED designed the study, conducted experiments, analyzed results, acquired funding and wrote the manuscript. ASC conducted experiments and analyzed results. JD, DLH and PR conducted experiments and provided scientific advice. EJC supervised data analysis and provided scientific advice. MMH and AL supervised research and edited the manuscript; MAL designed the study, supervised research, acquired funding and wrote the manuscript. All authors read, edited and approved the manuscript.

