## Supplementary material for "Intrinsic defects in lymph node stromal cells underpin poor germinal center responses during aging": Figure S1

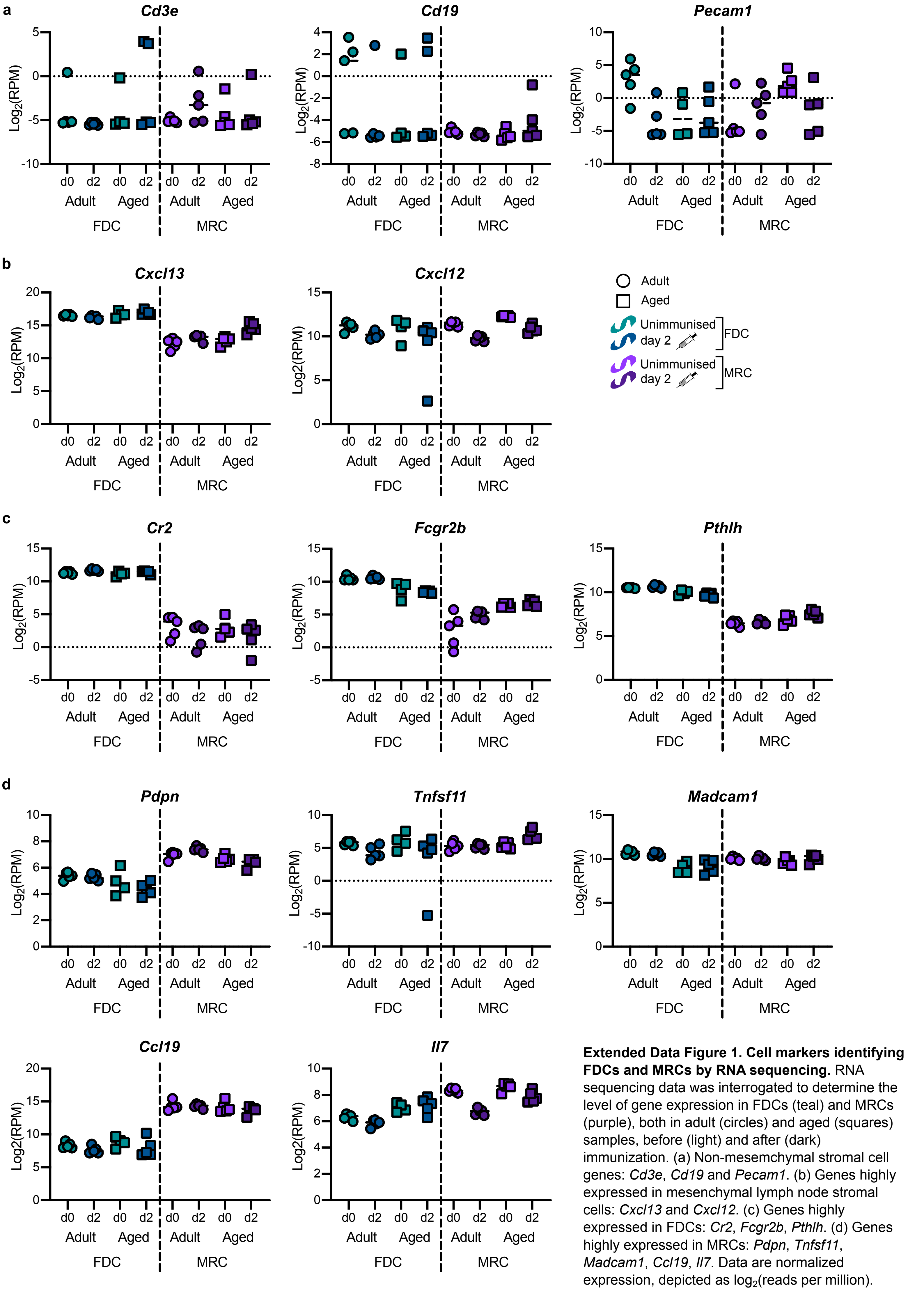


**Extended Data Figure 1. Cell markers identifying FDCs and MRCs by RNA sequencing.** RNA sequencing data was interrogated to determine the level of gene expression in FDCs (teal) and MRCs (purple), both in adult (circles) and aged (squares) samples, before (light) and after (dark) immunization. (a) Non-mesemchymal stromal cell genes: *Cd3e*, *Cd19* and *Pecam1*. (b) Genes highly expressed in mesenchymal lymph node stromal cells: *Cxcl13* and *Cxcl12*. (c) Genes highly expressed in FDCs: *Cr2*, *Fcgr2b*, *Pthlh*. (d) Genes highly expressed in MRCs: *Pdpn*, *Tnfsf11*, *Madcam1*, *Ccl19*, *Il7*. Data are normalized expression, depicted as $\log_2$(reads per million).
