## Supplementary material for "Intrinsic defects in lymph node stromal cells underpin poor germinal center responses during aging": Figure S3

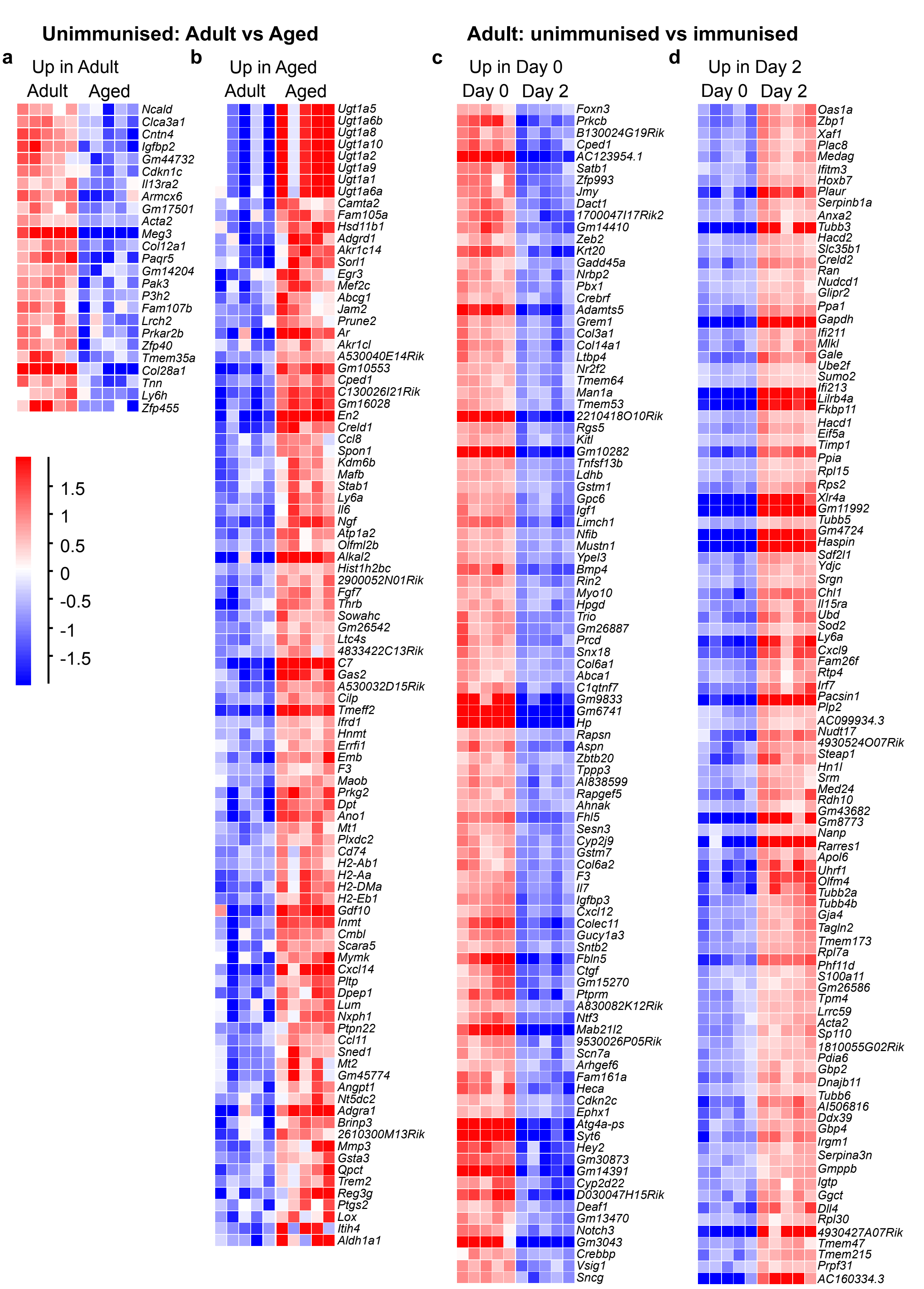


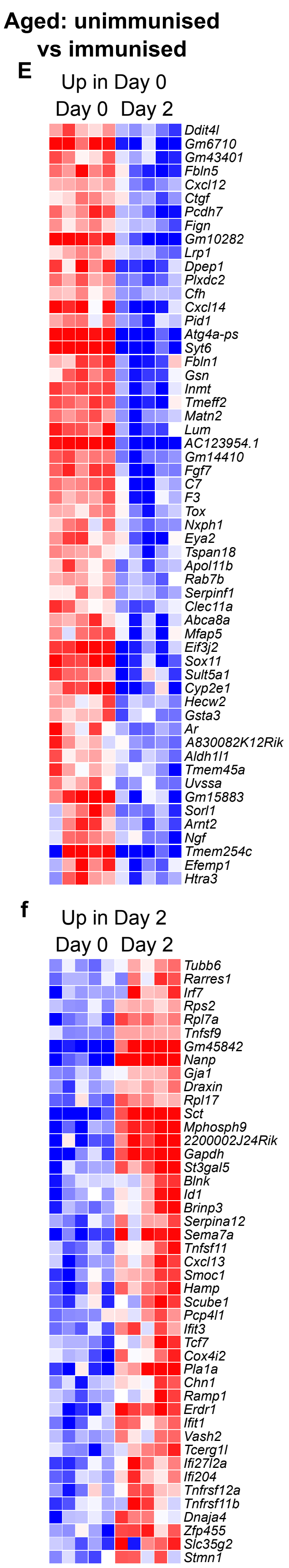


**Extended Data Figure 3. Differentially expressed genes in MRCs as determined by RNA sequencing.** Differentially expressed genes were determined in pairwise comparisons between adult and aged MRCs in the resting state (a, b) and prior to and following immunization in adult (c, d) and aged (e, f) MRCs. Differentially expressed genes were determined by DESeq2 on all expressed genes with a $\log_2$(fold change)>1 and an adjusted p-value < 0.05. Data are normalized within each gene; scale indicates normalized expression ($\log_2$).
