## Supplementary material for "Intrinsic defects in lymph node stromal cells underpin poor germinal center responses during aging": Figure S4

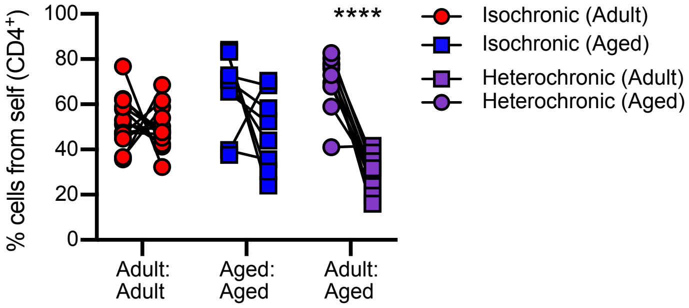


**Extended Data Figure 4. T cell chimerism in parabionts.** The proportion of T cells that were derived from the original host, based on CD45.1 or CD45.2 expression, was determined in blood for adult and aged isochronic parabionts and heterochronic parabionts. Statistical significance was determined using a two-way ANOVA with Sidak's multiple comparison test, ****p<0.0001.
